# The inhaled steroid ciclesonide blocks SARS-CoV-2 RNA replication by targeting viral replication-transcription complex in culture cells

**DOI:** 10.1101/2020.08.22.258459

**Authors:** Shutoku Matsuyama, Miyuki Kawase, Naganori Nao, Kazuya Shirato, Makoto Ujike, Wataru Kamitani, Masayuki Shimojima, Shuetsu Fukushi

## Abstract

We screened steroid compounds to obtain a drug expected to block host inflammatory responses and MERS-CoV replication. Ciclesonide, an inhaled corticosteroid, suppressed replication of MERS-CoV and other coronaviruses, including SARS-CoV-2, the cause of COVID-19, in cultured cells. The effective concentration (EC_90_) of ciclesonide for SARS-CoV-2 in differentiated human bronchial tracheal epithelial cells was 0.55 μM. Ciclesonide inhibited formation of double membrane vesicles, which anchor the viral replication-transcription complex in cells. Eight consecutive passages of 43 SARS-CoV-2 isolates in the presence of ciclesonide generated 15 resistant mutants harboring single amino acid substitutions in non-structural protein 3 (nsp3) or nsp4. Of note, ciclesonide still suppressed replication of all these mutants by 90% or more, suggesting that these mutants cannot completely overcome ciclesonide blockade. These observations indicate that the suppressive effect of ciclesonide on viral replication is specific to coronaviruses, highlighting it as a candidate drug for the treatment of COVID-19 patients.

**Importance:** The outbreak of SARS-CoV-2, the cause of COVID-19, is ongoing. To identify the effective antiviral agents to combat the disease is urgently needed. In the present study, we found that an inhaled corticosteroid, ciclesonide suppresses replication of coronaviruses, including beta-coronaviruses (MHV-2, MERS-CoV, SARS-CoV, and SARS-CoV-2) and an alpha-coronavirus (HCoV-229E) in cultured cells. The inhaled ciclesonide is safe; indeed, it can be administered to infants at high concentrations. Thus, ciclesonide is expected to be a broad-spectrum antiviral drug that is effective against many members of the coronavirus family. It could be prescribed for the treatment of MERS, and COVID-19.

## Introduction

The COVID-19 outbreak began in December 2019 in Wuhan, China(1). The causative virus, severe acute respiratory syndrome coronavirus 2 (SARS-CoV-2), spread rapidly worldwide and was declared a global health emergency by the World Health Organization. Thus, effective antiviral agents to combat the disease are urgently needed. Several drugs are effective against SARS-CoV-2 in cultured cells(2–4). Of these, Remdesivir has undergone clinical trials in COVID-19 patients, with both positive and negative results (5, 6). Lopinavir/ritonavir and chloroquine/hydroxychloroquine are of no benefit (7, 8).

The virus can have inflammatory effects; therefore, steroids are used to treat severe inflammation, with beneficial effects in some cases. For example, high-dose steroids reduce symptoms in those with influenza encephalopathy(9). It would be highly beneficial if a virus-specific inhibitor was identified among the many steroid compounds that have been well characterized. However, systemic treatment with steroids is contraindicated in cases of severe pneumonia caused by Middle East respiratory syndrome coronavirus (MERS-CoV) or severe acute respiratory syndrome coronavirus (SARS-CoV); this is because steroids suppress innate and adaptive immune responses(10, 11), resulting in increased viral replication. In fact, for SARS (2003) and MERS (2013), systemic treatment with cortisone or prednisolone is associated with increased mortality(12, 13). Therefore, if steroid compounds are to be used to treat patients suffering from COVID-19, their tendency to increase virus replication must be abrogated. Here, we reconsidered the use of steroids for treatment of pneumonia caused by coronavirus.

In the preprint of this study (posted in BioRxiv)(14), we showed that a corticosteroid, ciclesonide, is a potent blocker of SARS-CoV-2 replication. Based on the data in our preprint study, clinical trials of a retrospective cohort study to treat COVID-19 patients were started in Japanese hospital in March 2020. The treatment regime involves inhalation of 400 μg ciclesonide (two or three times per day) for 2 weeks. Three cases of COVID-19 pneumonia treated successfully with ciclesonide have been reported(15), as have several case reports(16–18). None of these studies reported significant side effects. The aim of the present study is to outline the scientific rationale for conducting these clinical trials.

## Results

### Antiviral effect of steroid compounds on MERS-CoV

The 92 steroid compounds chosen from the Prestwick Chemical Library were examined to assess the inhibitory effects of MERS-CoV-induced cytopathic effects. Vero cells treated with steroid compounds were infected with MERS-CoV at an MOI = 0.1 and then incubated for 3 days. Four steroid compounds, ciclesonide, mometasone furoate, mifepristone, and algestone acetophenide, conferred a > 95% cell survival rate (Fig. 1). Interestingly, a structural feature of these compounds is a five- or six-membered monocycle attached to the steroid core.

**Figure 1.**
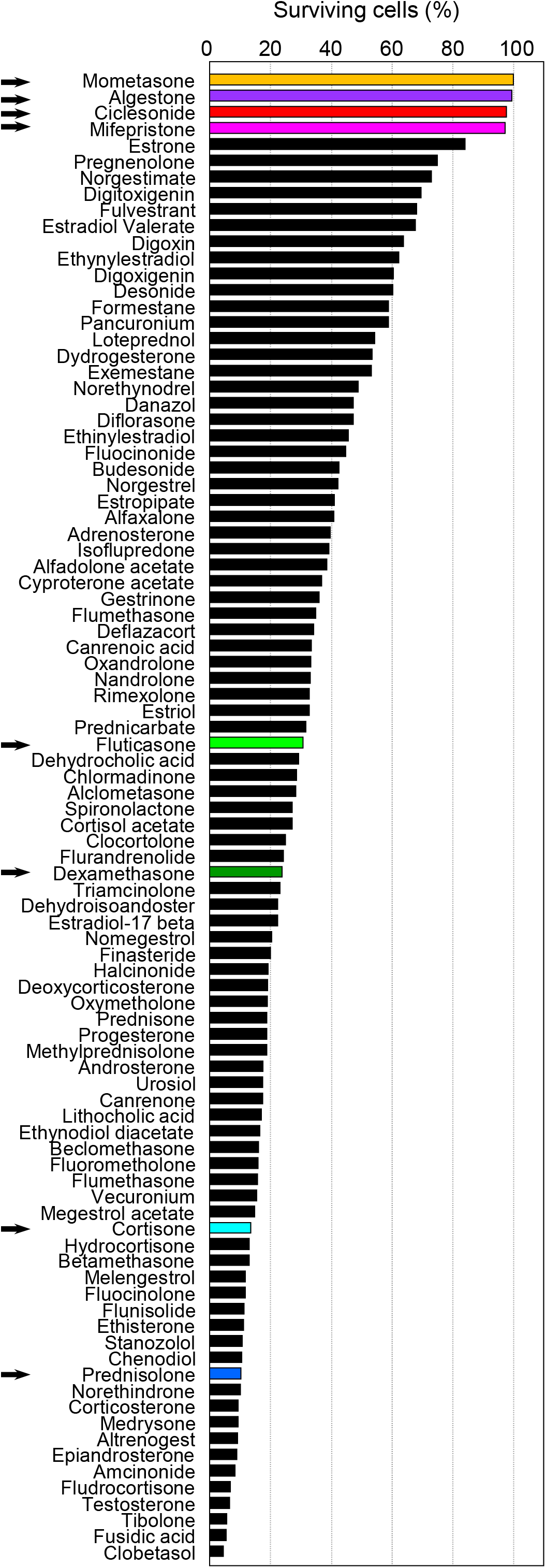
Steroid compounds reduce death rate of MERS-CoV infected cells. Vero cells seeded in 96-well microplates were infected with 100 TCID_50_ MERS-CoV in the presence of steroid compounds (10 μM). Cytopathic effects were observed at 72 h post-infection. Surviving cells were stained with crystal violet and photographed and quantified using ImageJ software. Data are presented as the average of two independent wells. Arrows indicate the steroid compounds assessed further in this study.

Next, we assessed the ability of eight steroid compounds (denoted by arrows in Fig. 1) to suppress both growth of MERS-CoV and virus-mediated cytotoxicity in Vero cells over a range of drug concentrations (0.1–100 µM). Ciclesonide exhibited low cytotoxicity and potent suppression of viral growth (Fig. 2a). Algestone acetophenide, mometasone, and mifepristone also suppressed viral growth; however, at 10 μM, the percentage viability of cells treated with algestone acetophenide and mometasone was lower than that of cells treated with ciclesonide, and the ability of mifepristone and mometasone to suppress viral growth was lower than that of ciclesonide. Cortisone and prednisolone, which are commonly used for systemic steroid therapy, dexamethasone, which has strong immunosuppressant effects, and fluticasone, a common inhaled steroid drug, did not suppress viral growth (Fig. 2a). A time-of-addition assay to compare the viral inhibition efficacies of the steroids with those of E64d, a cathepsin-dependent virus entry inhibitor, and lopinavir, a viral 3CL protease inhibitor, previously reported for SARS-CoV(19, 20), demonstrated that ciclesonide functions at the post-virus entry stage (Supplemental Fig. S1).

**Figure 2.**
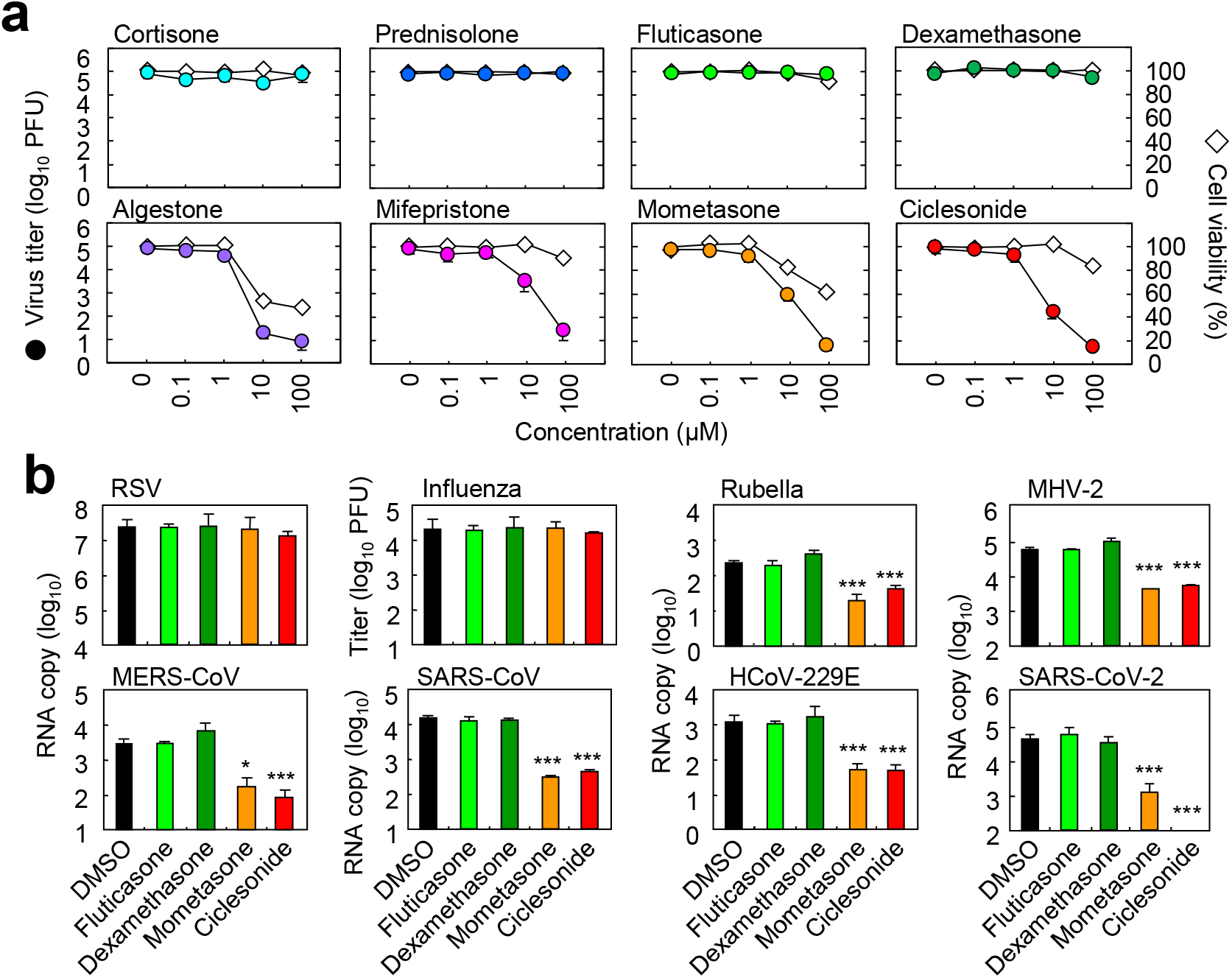
Steroid compounds suppress replication of MERS-CoV and other viruses. **(a)** The effects of eight steroid compounds on MERS-CoV replication. Vero cells were infected with MERS-CoV at an MOI of 0.01 in the presence of the indicated steroids for 24 h. The viral titer in the cell supernatant was quantified in a plaque assay using Vero/*TMPRSS2* cells. Cell viability in the absence of virus was quantified in a WST assay. **(b)** The antiviral effects of steroid compounds on various viral species. Cells were infected with the indicated viruses at an MOI of 0.01 in the presence of DMSO (control) or the indicated steroids. The viral yield in the cell supernatant was quantified in plaque assay or real-time PCR. Hep-2 cells were incubated with respiratory syncytial virus A (RSV-A long) for 1 day; MDCK cells were incubated with influenza H3N2 for 1 day; Vero cells were incubated with rubella virus (TO336) for 7 days; DBT cells were incubated with murine coronavirus (MHV-2) for 1 day; Vero cells were incubated with MERS-CoV (EMC), SARS-CoV (Frankfurt-1), or SARS-CoV-2 (WK-521) for 1 day; and HeLa229 cells were incubated with HCoV-229E (VR-740) for 1 day.

The antiviral effects of mometasone and ciclesonide against various viral species were tested by quantifying propagated virus in the culture medium of infected cells. Ciclesonide and mometasone suppressed replication of MHV-2, MERS-CoV, SARS-CoV, HCoV-229E, and SARS-CoV-2 (all of which have a positive strand RNA genome), but did not affect replication of respiratory syncytial virus (RSV) or influenza virus (which have a negative strand RNA genome) (Fig. 2b). In addition, ciclesonide slightly, but significantly, inhibited replication of rubella virus (which has a positive strand RNA genome) (Fig. 2b).

### Target of ciclesonide during MERS-CoV replication

In an attempt to identify a druggable target for viral replication, we performed 11 consecutive passages of MERS-CoV in the presence of 40 μM ciclesonide or 40 μM mometasone. A mutant virus displaying resistance to ciclesonide (but no virus displaying resistance to mometasone) was generated. Viral replication in the presence of ciclesonide was confirmed by measuring the virus titer in the culture medium of infected Vero cells at 24 h post-infection (hpi) and the amount of viral RNA in infected cells at 6 hpi (Fig. 3a and 3b). Next-generation sequencing revealed that an amino acid substitution at A25V (C19647T in the reference sequence NC_019843.3) in non-structural protein 15 (NSP15), a coronavirus endoribonuclease (21–23), was predicted to cause resistance to ciclesonide. Subsequently, a recombinant virus carrying the A25V amino acid substitution in nsp15 (Re-Nsp15-A25V) was generated from the parental MERS-CoV/EMC strain (Re-EMC/MERS) using a bacterial artificial chromosome (BAC) reverse genetics system(24). The titer of recombinant virus in the culture medium of infected Vero cells at 24 hpi, and the amount of viral RNA in the cells at 6 hpi, were quantified. As expected, the Re-Nsp15-A25V strain was much less susceptible to ciclesonide than the parental strain (Fig. 3c and 3d).

**Figure 3.**
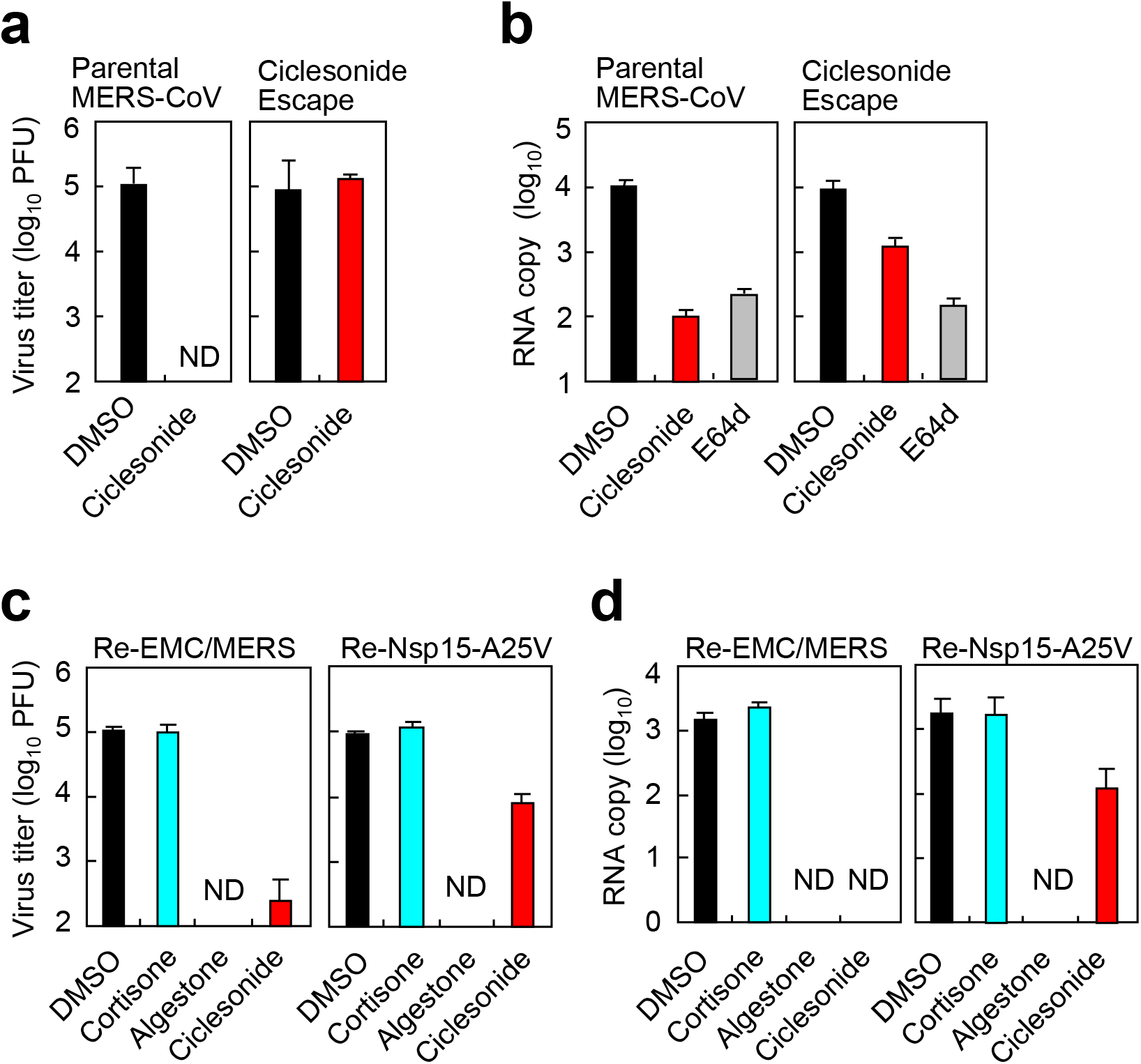
A ciclesonide-escape mutant of MERS-CoV. **(a)** Viral growth of a ciclesonide escape MERS-CoV mutant. Vero cells treated with 10 µM ciclesonide were infected with parental MERS-CoV or the ciclesonide escape mutant at an MOI of 0.01. The viral titer in the culture medium was quantified at 24 h post-infection (hpi). **(b)** Viral RNA replication of a ciclesonide escape MERS-CoV mutant. Vero cells treated with 10 µM ciclesonide were infected with parental MERS-CoV or the ciclesonide escape mutant at an MOI of 1. The viral RNA in the cells was quantified at 6 hpi. E64d (10 µM), a virus entry inhibitor, was used for comparison. **(c)** Growth of the recombinant virus. Vero cells were infected with the parental MERS-CoV/EMC strain (Re-EMC/MERS) or the recombinant mutant strain (Re-Nsp15-A25V) containing an amino acid substitution at A25V in NSP15 at an MOI of 0.01 and then treated with the indicated compounds (10□μM). Virus titer was quantified at 24 hpi. **(d)** RNA replication of the recombinant virus. Vero cells were infected with Re-EMC/MERS or Re-Nsp15-A25V at an MOI of 1 and treated with the indicated compounds (10□μM). The viral RNA in infected cells was quantified at 6 hpi. ND, not detected. Data are represented as the mean ± standard deviation of four independent wells; * P ≤ 0.05; and *** P ≤ 0.001.

### Antiviral effect of steroid compounds on SARS-CoV-2

In response to the global outbreak of COVID-19, our study target changed from MERS-CoV to SARS-CoV-2. We evaluated the inhibitory effects of ciclesonide on replication of the latter. First, the effective concentration of ciclesonide required to inhibit virus propagation was assessed by quantifying the virus titer in the supernatant of VeroE6/*TMPRSS2* cells at 24 hpi (Fig. 4a and 4b); this cell line is highly susceptible to SARS-CoV-2(25). We also examined human bronchial epithelial Calu-3 cells (Fig. 4c and 4d). Ciclesonide blocked SARS-CoV-2 replication in a concentration-dependent manner (EC_90_□= □LJ5.1□μM in VeroE6/*TMPRSS2* cells; EC_90_□= □6.0□μM in Calu-3 cells). In addition, differentiated primary human bronchial tracheal epithelial (HBTE) cells at an air-liquid interface (ALI) (HBTE/ALI cells) were prepared and SARS-CoV-2 replication was evaluated in the presence of ciclesonide. At 3 days post-infection, real-time PCR revealed a 2,000-fold increase in the amount of viral RNA in cells (Fig. 4e); ciclesonide suppressed replication of viral RNA at a low concentration (Fig. 4f) (EC_90_□= □0.55□μM in HBTE/ALI cells). The amount of viral RNA detected in the liquid phase was low, indicating that less virus is secreted via the basolateral surface (Fig. 4f). To assess the effect of ciclesonide at the early stage of SARS-CoV-2 replication, we measured the amount of viral RNA in VeroE6/*TMPRSS2* cells over time. The quantitative level of RNA replication was observed at 6 h post-infection (Fig. 5a). Nelfinavir and lopinavir, strong inhibitors of SARS-CoV-2 RNA replication (4, 26), were used for comparison. At 6 hpi with SARS-CoV-2 (MOI = 1), mometasone and ciclesonide suppressed viral RNA replication with an efficacy similar to that of nelfinavir and lopinavir; however, fluticasone and dexamethasone did not suppress replication (Fig. 5b).

**Figure 4.**
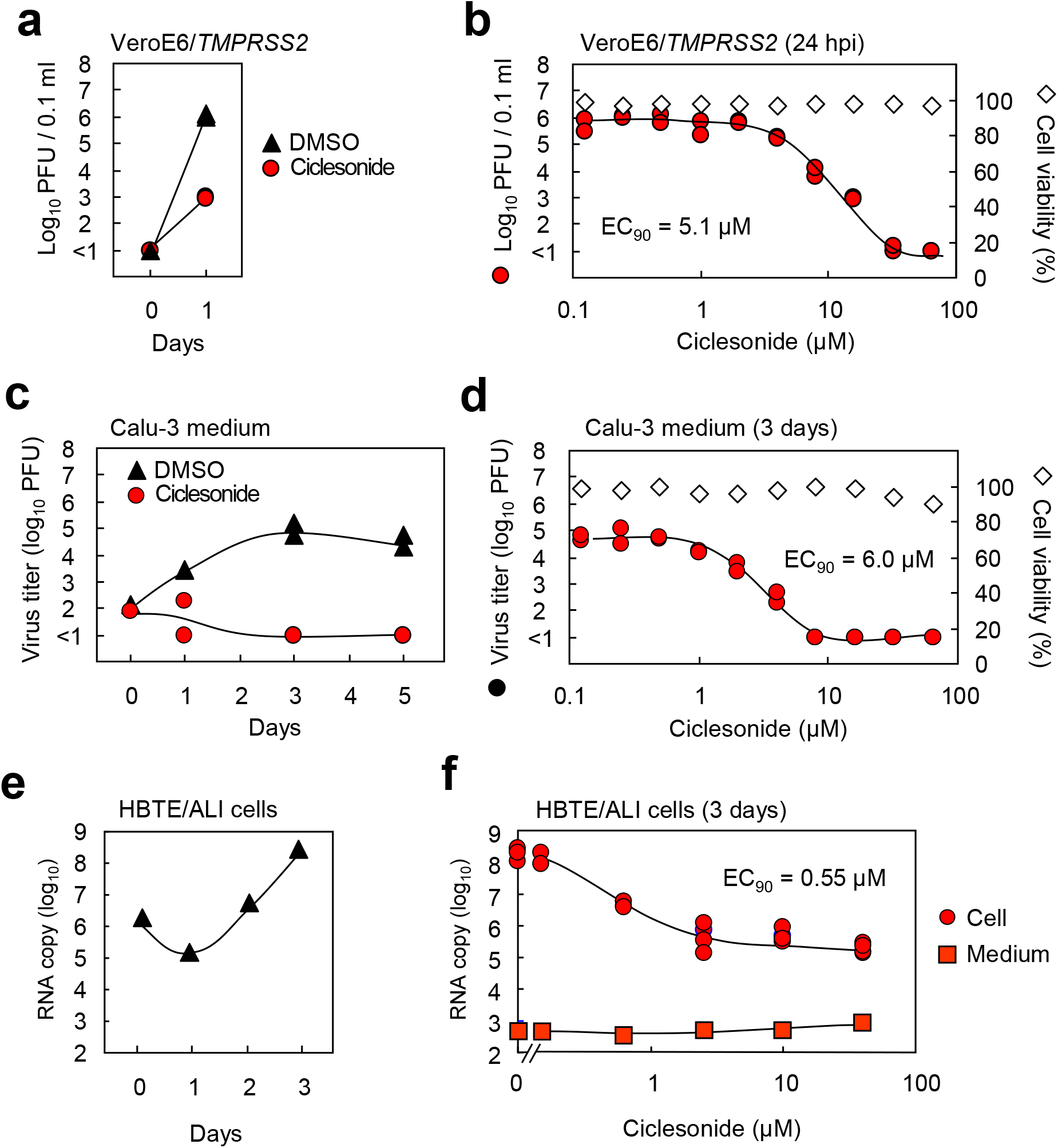
Ciclesonide suppresses replication of SARS-CoV-2 in human bronchial cells. **(a, c** and **e)** Time course of SARS-CoV-2 propagation. **(b, d** and **f)** Concentration-dependent effects of ciclesonide. VeroE6/*TMPRSS2* cells (panels a and b), Calu-3 cells (panels c and d) or HBTE/ALI cells (panels e and f) were infected with SARS-CoV-2 at an MOI of 0.001 in the presence of DMSO or ciclesonide (10 μM) and then incubated for 1, 3 or 5 days. The virus titer in medium was quantified in a plaque assay using VeroE6/*TMPRSS2* cells (n=2, in panels a and c); alternatively, the viral RNA in cells or culture medium was quantified by real-time PCR using the E gene primer/probe set (n=1 in panel e, or n=4 in panel f). Average of cell viability in the absence of virus was quantified using a WST assay (n=2, in panel b and d).

**Figure 5.**
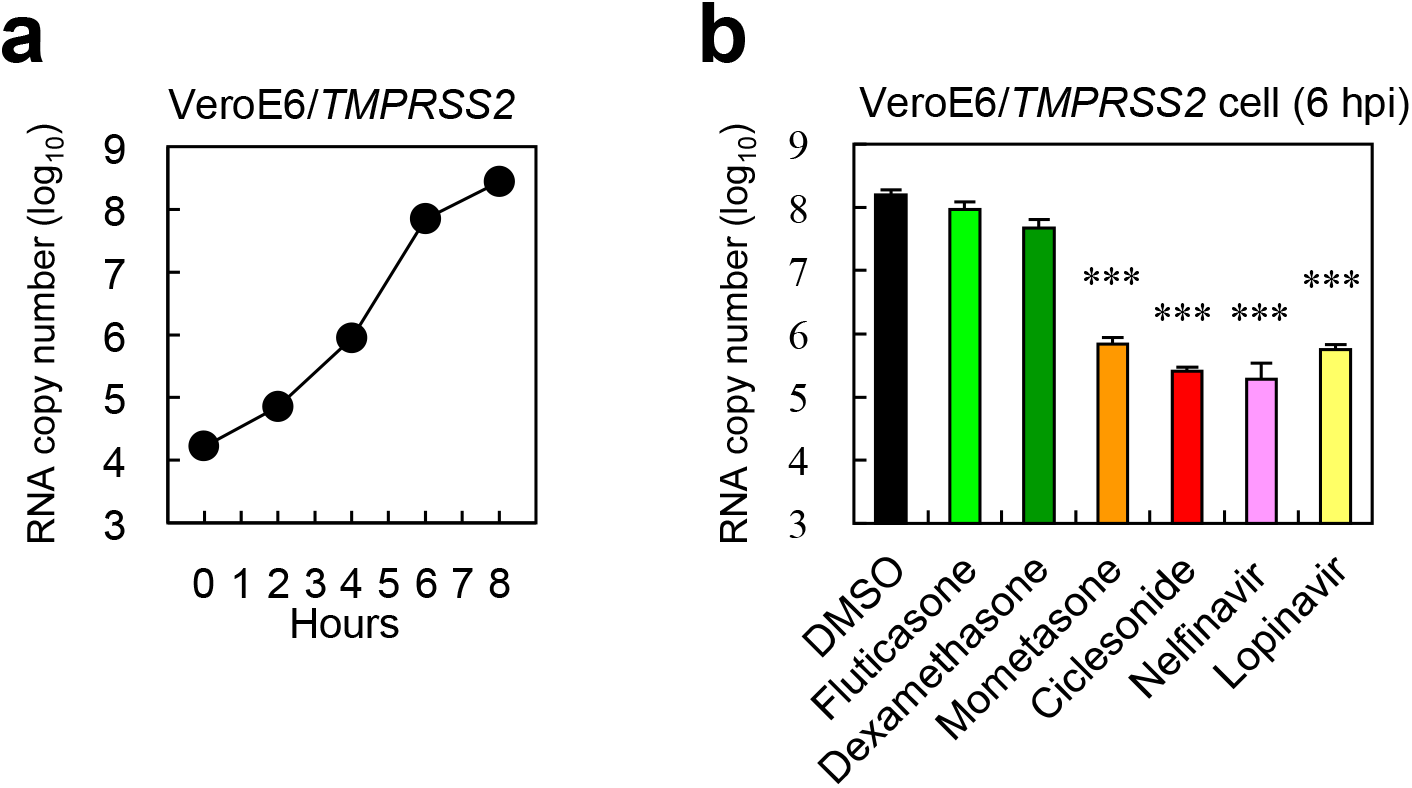
Steroid compounds and other inhibitors suppress SARS-CoV-2 RNA replication in VeroE6/*TMPRSS2* cells. **(a)** Time course of SARS-CoV-2 RNA replication. Cells were infected with virus at an MOI of 1 and cellular RNA was collected at the indicated time points. **(b)** Inhibition of viral RNA replication. Cells were infected with SARS-CoV-2 at an MOI of 1 in the presence of the indicated compounds (10□μM) for 6 h. Cellular viral RNA was quantified by real-time PCR using the E gene primer/probe set. *** P ≤ 0.001.

### Target of ciclesonide during SARS-CoV-2 replication

To identify the molecule targeted by ciclesonide to suppress viral RNA replication, we used 43 SARS-CoV-2 isolates from infected patients to generate ciclesonide escape mutants. Consecutive passage of these isolates in VeroE6/*TMPRSS2* cells in the presence of 40 μM ciclesonide. After eight passages, three viral plaques from each passage of the 43 cell supernatants were isolated in a limiting dilution assay; the viral RNA was then isolated for next-generation sequencing. We obtained 15 isolates harboring a single mutation in the viral genome when compared with that of the parental virus (Table 1). We examined replication of these mutants in the presence of ciclesonide. First, one of these isolates was tested in VeroE6/*TMPRSS2* cells. At 6 hpi, the amount of viral RNA derived from the parental virus fell by 1000-fold in the presence of ciclesonide; by contrast, the amount of RNA derived from the escape mutant increased 50-fold compared with that of the parent virus (Fig. S2). There was no difference between the parental virus and the escape mutant in the presence of other steroid compounds (i.e., cortisone and algestone acetophenide) (Fig. S2). Furthermore, when we tested all 15 mutants in the presence of ciclesonide, we found a 6-to 50-fold increase in the amount of mutant viral RNA compared with that of the parental virus (Fig. 6a). Importantly, ciclesonide suppressed replication of all escape mutants by 90% or more, suggesting that these mutants cannot completely overcome ciclesonide blockade. Mutations in the ciclesonide escape mutants were identified at three positions in nsp3 and at one position in nsp4 (Fig. 6b). Of note, the amino acid substitution N1543K in nsp3 was caused by a different base change (T7348G and T7348A) (Table 1). Nsp3 and nsp4 are involved in formation of double membrane vesicles (DMV), which anchor the coronavirus replication-transcription complex within cells (27, 28). In VeroE6/*TMPRSS2* cells, DMVs were observed at 5 hpi using an anti-SARS-CoV nsp3 antibody and an anti-double strand RNA antibody (Fig. 7). The fluorescence intensity of these molecules fell in the presence of ciclesonide in a concentration-dependent manner (Fig. 7).

**Table 1.** Mutation of ciclesonide escape mutant

| Coronavirus species and reference sequence | Mutation in viral genome | Amino acid position in ORF1a | Amino acid position in nsp | passage number | Parental strain of mutant |
| --- | --- | --- | --- | --- | --- |
| MERS-CoV-2, NC019843.3 | C19647T | 6457 | nsp15, A25V | 11 | EMC/2012 |
| SARS-CoV-2, MN908947.3 | T7348G | 2361 | nsp3, N1543K | 8 | DP15-134, DP15-200, DP16-090, DP16-281, DP17-144 |
|  | T7348A | 2361 | nsp3, N1543K | 8 | WK-521 |
|  | G8006A | 2581 | nsp3, G1763S | 8 | DP15-078, DP15-196, DP16-074, DP16-157, DP16-282, DP17-187 |
|  | A8010C | 2582 | nsp3, D1764A | 8 | DP17-243 |
|  | G9242A | 2994 | nsp4, E230K | 8 | DP15-104, DP16-238 |

**Figure 6.**
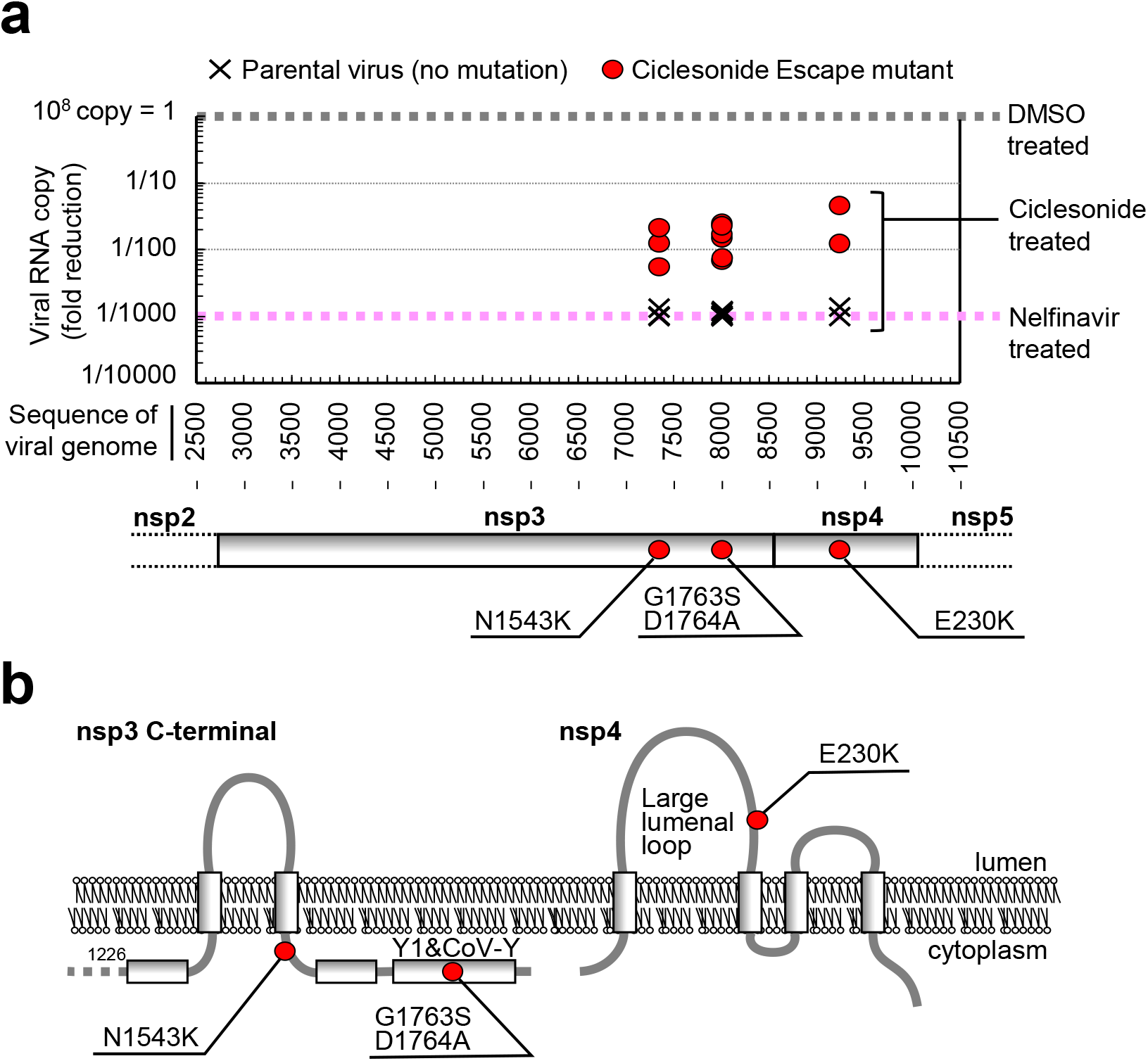
A ciclesonide escape mutant of SARS-CoV-2. **(a)** Virus replication in the presence of ciclesonide is due to amino acid substitutions in nsp3 and nsp4. Replication of RNA derived from the 15 mutants listed in Table 1 was assessed in VeroE6/*TMPRSS2* cells. Viral RNA was isolated at 6 hpi and measured by real-time PCR using the E gene primer/probe set. The results were compared with those for the parental virus in which the viral RNA level after treatment with DMSO was set to 1, and that after treatment with nelfinavir was set to 1/1000. Relative reductions of viral RNA in the presence of ciclesonide were plotted at the corresponding mutations in the SARS-CoV-2 genome sequence. The amino acid substitutions in nsp3 and nsp4 are shown at the bottom of the panel. **(b)** Topological diagram. The C-terminal region of nsp3 and full-length nsp4 are depicted on the lipid bilayer of the endoplasmic reticulum membrane.

**Figure 7.**
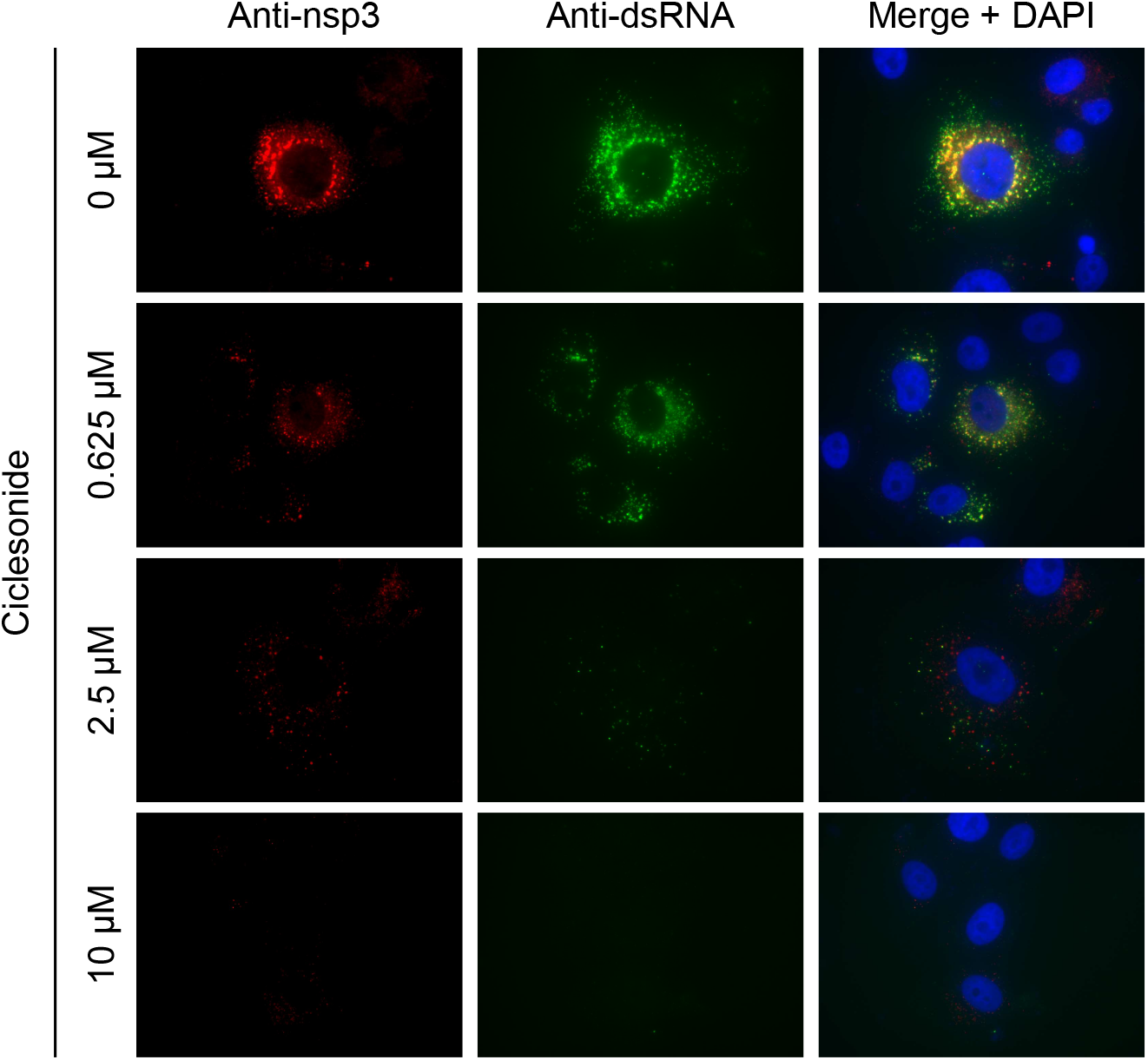
Ciclesonide suppresses DMV formation. VeroE6/*TMPRSS2* cells were infected with SARS-CoV-2 at an MOI of 0.1 in the presence of DMSO or ciclesonide, and then incubated for 5 h. Next, cells were fixed with 4% paraformaldehyde and permeabilized with 0.1% Tween-20. Nsp3 and double strand RNA were stained with a rabbit anti-SARS-nsp4 antibody and a mouse anti-dsRNA antibody, followed by Alexa Fluor 594 conjugated anti-rabbit IgG and Alexa Fluor 488-conjugated anti-mouse IgG. Cell nuclei were stained with DAPI.

## Discussion

Inhaled ciclesonide is safe; indeed, it can be administered to infants at high concentrations. Because it remains primarily in the lung tissue and does not enter the bloodstream to any significant degree(29), its immunosuppressive effects are weaker than those of cortisone and prednisolone(29, 30). The data presented herein suggest that inhaled ciclesonide has the potential to reduce both viral replication and inflammation in the lungs.

We found that ciclesonide suppresses replication of coronaviruses, including beta-coronaviruses (MHV-2, MERS-CoV, SARS-CoV, and SARS-CoV-2) and alpha-coronaviruses (HCoV-229E) in cultured cells. Thus, ciclesonide is expected to be a broad-spectrum antiviral drug that is effective against many members of the coronavirus family. It could be prescribed for the treatment of common colds, MERS, and COVID-19. The concentration of ciclesonide that effectively reduced replication of SARS-CoV-2 in differentiated HBTE cells was 10-fold lower than that required to suppress replication in VeroE6/*TMPRSS2* or Calu-3 cells (Fig. 4b, 4d and 4f). It is speculated that ciclesonide is a prodrug that is metabolized in lung tissue to yield the active form (31); therefore, it may be converted into its active form in differentiated HBTE cells. Furthermore, this study predicts the occurrence of ciclesonide escape mutants in patients treated with ciclesonide; however, the drug suppresses replication of these mutants by >90% (Fig. 6a). Until now, the mutations identified in these mutants have not been detected in SARS-CoV-2 sequences posted in the GISAID and NCBI databases.

A ciclesonide escape mutant of MERS-CoV harbored an amino acid substitution at the dimerization site of the NSP15 homo-hexamer(32)(32)(32). Nsp15 is an uridylate-specific endoribonuclease, an RNA endonuclease, which plays a critical role in coronavirus replication(32, 33). Recently, an *in silico* study suggested direct interaction between ciclesonide and nsp15 of SARS-CoV-2(34). However, we did not identify mutations in nsp15 of the ciclesonide escape mutants of SARS-CoV-2; rather, we identified mutations in the C-terminal cytosolic region (next to the transmembrane domain or within the Y1&CoV-Y domain) of nsp3 or in the large lumenal loop of nsp4 (Fig. 6b). Nsp3 contains a papain-like protease and, due to the large number of interactions with other nsps (including nsp4 and nsp15), is believed to be part of the central scaffolding protein of the replication-transcription complex (33, 35, 36). In addition, like coronavirus DMV, the rubella virus (which has a positive strand RNA genome and forms a spherule-like structure in cells) was slightly suppressed by ciclesonide, suggesting that ciclesonide may interact with the replication-transcription complex, a structure common to rubella and coronavirus. It is difficult to identify the mechanism by which ciclesonide targets the nsps complex because little is known about the construction and interaction of nsps in the replication-transcription complex. The result in Figure 7 was unable to be ascertain whether DMV formation or RNA replication was inhibited first by ciclesonide. We anticipate that further experiments using mutant nsps may reveal the molecular mechanism underlying the antiviral effect of ciclesonide.

## Materials and Methods

### Cells and viruses

Hep-2, HeLa229, MDCK, Calu-3, Vero, Vero/*TMPRSS2* and VeroE6/*TMPRSS2* cells were maintained in Dulbecco’s modified Eagle medium high glucose (DMEM, Sigma-Aldrich, USA), and DBT cells were maintained in DMEM (Nissui, Japan), supplemented with 5% fetal bovine serum (Gibco-BRL, USA). MERS-CoV and SARS-CoV-2 were propagated in Vero and VeroE6/*TMPRSS2* cells, respectively. HBTE cells (KH-4099; Lifeline cell technology, USA) were plated on 6.5-mm-diameter Transwell permeable supports (3470; Corning, USA), and human airway epithelium cultures were generated by growing the cells at an air-liquid interface for 3 weeks, resulting in well-differentiated, polarized cultures. For treatment of HBTE cells in the experiments, ciclesonide was mixed in liquid phase medium at the indicated concentrations and virus was inoculated onto the air-phase.

### Steroids and inhibitors

The following compounds were used: cortisone, prednisolone, fluticasone, dexamethasone, algestone acetophenide, mifepristone, mometasone furoate, ciclesonide (all from the Prestwick Chemical Library; PerkinElmer, USA), E64d (330005; Calbiochem, USA); nelfinavir (B1122; ApexBio, USA); and lopinavir (SML1222; Sigma-Aldrich).

### Quantification of viral RNA

Confluent cells in 96-well plates were inoculated with virus in the presence of steroid compounds. Cellular RNA was isolated at 6 hpi using the CellAmp Direct RNA Prep Kit (3732; Takara, Japan). The RNA was then diluted in water and boiled. Culture medium was collected at the indicated time points, diluted 10-fold in water, and then boiled. Real-time PCR assays to measure the amount of coronavirus RNA were performed using a MyGo Pro instrument (IT-IS Life Science, Ireland). The primers and probes are described in Supplemental Table S1. Viral mRNA levels were normalized to the expression levels of the cellular housekeeping gene GAPDH.

### Cytotoxicity Assays

Confluent cells in 96-well plates were treated with steroid compounds. After incubation for 24 or 27 h, a cell viability assay was performed using WST reagent (CK12; Dojin Lab, Japan), according to the manufacturer’s instructions.

### Generation of recombinant MERS-CoV from BAC plasmids

A BAC clone carrying the full-length infectious genome of the MERS-CoV EMC2012 strain, pBAC-MERS-wt, was used to generate recombinant MERS-CoV, as described previously(24, 37) The BAC DNA of SARS-CoV-Rep (38), kindly provided by Luis Enjuanes, was used as a backbone BAC sequence to generate pBAC-MERS-wt. The BAC infectious clones carrying amino acid substitutions in nsp15 was generated by modification of the pBAC-MERS-wt (as a template) using a Red/ET Recombination System Counter-Selection BAC Modification Kit (Gene Bridges, Heidelberg, Germany). BHK-21 cells were grown in a single well of a six-well plate in 10% FCS-MEM and transfected with 3 μg of BAC plasmid with Lipofectamine 3,000 (Thermo Fisher, USA). After transfection, Vero/TMPRSS2 cells were inoculated to transfected BHK-21 cells. The co-culture was then incubated at 37°C for 3 days. The supernatants were collected and propagated once using Vero/TMPRSS2 cells. Recovered viruses were stored at −80°C.

### Generation of ciclesonide escape mutant

To obtain ciclesonide escape mutants, virus passage was repeated at least eight times in the presence of 40 μM ciclesonide. At the first passage, about 10^7^ PFU of virus was inoculated onto 10^6^ cells and incubated for 3 h. Next, the cells were washed twice with culture medium and incubated for 2 days in the presence of ciclesonide. The incubation period was 2 days for the first three passages and 1 day for the following passages. Cells were inoculated with 100 μl culture medium at each successive passage. The amount of replicating virus in the presence of ciclesonide was quantified using real-time PCR. Vero and VeroE6/*TMPSS2* cells were used to passage MERS-CoV and SARS-CoV-2, respectively.

### Whole genome sequencing of SARS-CoV-2

Extracted viral RNA was reverse transcribed and tagged with index adaptors using the NEBNext Ultra II RNA Library Prep Kit for Illumina (New England Biolabs, Ipswich, MA, USA), according to the manufacturer’s instructions. The resulting cDNA libraries were verified using the MultiNA System (Shimadzu, Kyoto, Japan) and quantified using a Quantus Fluorometer (Promega, Madison, WI, USA). Indexed libraries were then converted and sequenced (150-bp paired-end reads) using the DNBSEQ-G400 (MGI Tech., Shenzhen, China; operated by GENEWIZ, South Plainfield, NJ, USA). After sequencing, reads with the same index sequences were grouped. Sequence reads were trimmed by Ktrim (39) and mapped onto the viral genomes of parental strains using Minimap2 (40). The consensus sequences of the mapped reads were obtained using ConsensusFixer (Töpfer A. https://github.com/cbg-ethz/consensusfixer).

### Immunofluorescence Microscopy

VeroE6/*TMPRSS2* cells cultured on 96 well plates (Lumos multiwell 96; 94 6120 096; Sarstedt, Germany) were infected with SARS-CoV-2 (WK-521) at an MOI = 0.1 and incubated for 5 h. Next, the cells were fixed for 30 min at 4°C with 4% paraformaldehyde in phosphate-buffered saline (PBS). After washing once with PBS, the cells were permeabilized for 15 min at room temperature (RT) with PBS containing 0.1% Tween-20. The cells were then incubated with a mixture of rabbit anti-SARS-nsp4 (1:500; ab181620; Abcam, USA) and mouse anti-dsRNA (1:1000; J2-1709; Scicons, Hungary) antibodies for 1 h at RT, washed three times with PBS, and incubated for 1 h at RT with a mixture of Alexa Fluor 594 conjugated anti-rabbit IgG (1:500; A11012; ThermoFisher, USA) and Alexa Fluor 488-conjugated anti-mouse IgG (1:500; A10680; ThermoFisher, USA). Next, the cells were washed three times with PBS and cell nuclei were stained with DAPI (1:5000; D1306; ThermoFisher, USA). Cells were observed under an inverted fluorescence phase contrast microscope (BZ-X810; Keyence, Japan).

### Statistical analysis

Statistical significance was assessed using ANOVAs. P < 0.05 was considered statistically significant. In figures with error bars, data are presented as the mean ± SD.

## Acknowledgments

We are most grateful to Tsuneo Morishima (Aichi Medical University) for helpful suggestions. We also thank Yuriko Tomita, Makoto Kuroda, Tsuyoshi Sekizuka, Ikuyo Takayama, Mina Nakauchi, Kazuya Nakamura, Tsutomu Kageyama, Kazuhiko Kanou, Kiyoko Okamoto, Naoko Iwata, and Noriyo Nagata (National Institute of Infectious Diseases) for providing reagents and important information, Ron A. M. Fouchier and Bart L. Haagmans (Erasmus Medical Center) for providing MERS-CoV, and John Ziebuhr (University of Wurzburg) for providing SARS-CoV. This study was supported by Grants-in Aid from the Japan Agency for Medical Research and Development (AMED) (grant number JP19fk0108058j0802 and 20fk0108058j0803), and from the Japan Society for the Promotion of Science (JSPS) (grant number 17K08868 and 20K07519).

## Notes

https://www.biorxiv.org/content/10.1101/2020.03.11.987016v1

### Competing Interest Statement

The authors have declared no competing interest.

