## Supplemental Figure S1 for "The inhaled steroid ciclesonide blocks SARS-CoV-2 RNA replication by targeting viral replication-transcription complex in culture cells"

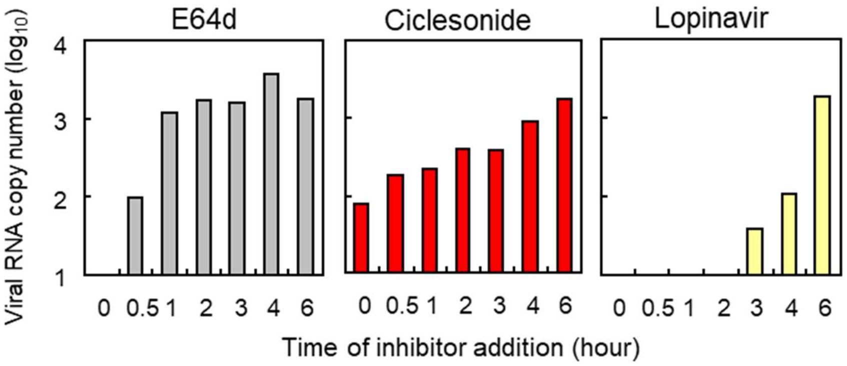


**Supplemental Fig. S1. Time-of-addition assays for MERS-CoV replication inhibitors.** The inhibitors E64d, ciclesonide, and lopinavir (each at 10 μM) were added to Vero cells at the indicated times after virus inoculation. The amount of cellular viral mRNA at 6 h post-infection was measured by real-time PCR.
