## Supplemental Figure S2 for "The inhaled steroid ciclesonide blocks SARS-CoV-2 RNA replication by targeting viral replication-transcription complex in culture cells"

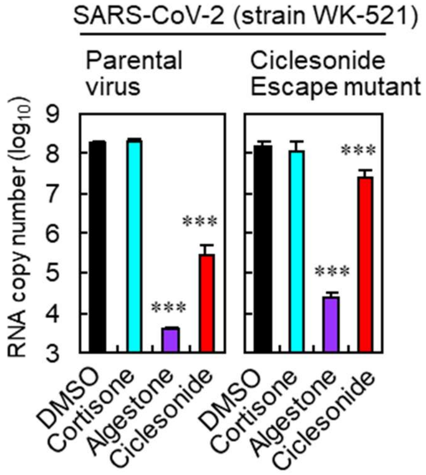


**Supplemental Fig. S2. A ciclesonide escape mutant of SARS-CoV-2.** VeroE6/*TMPRSS2* cells treated with the indicated compounds (each at 10 μM) were infected with parental SARS-CoV-2 or with the ciclesonide escape mutant (MOI = 1). Viral RNA titers in cells were measured at 6.5 hpi. Data are presented as the mean ± SD of n = 4 independent experiments; *** P ≤ 0.001.
